# Beneficial fungi are major drivers of root fungal microbiome assembly and network robustness

**DOI:** 10.1101/2025.04.27.650865

**Authors:** Saveria Mosca, Edda Francomano, Meriem Miyassa Aci, Nesma Zakaria Mohamed, Leonardo Schena, Antonino Malacrinò

## Abstract

Despite the critical role of soil microbiomes in ecosystem stability, how soil disturbance impacts these communities and their interactions with plants remains poorly understood. In this study, we investigate how different levels of soil disturbance influence the assembly and robustness of the root fungal microbiome. Using a microcosm experiment, we inoculated plants with soil from environments with decreasing levels of disturbance — agricultural field, field margin, and uncultivated field — and tested the variation in composition, assembly process, and network stability of the root fungal microbiome. Our results reveal that more disturbed systems, i.e. agricultural soils, harbor microbiomes with less robust networks compared to systems with lower disturbance. Network analysis identified arbuscular mycorrhizal fungi as key taxa contributing to microbiome stability, suggesting their critical role in maintaining the robustness of root-associated fungal microbiomes. Furthermore, stochastic processes dominated fungal community assembly across all soil treatments, yet the key taxa that deviated from null-model predictions include those with a major role in network robustness, and were mainly identified as arbuscular mycorrhizal fungi. Together, our results suggest that arbuscular mycorrhizal fungi are major actors in assembling robust plant microbiomes, and soil disturbance is a key factor in disrupting these interactions.

## Introduction

Global soils are currently under the threat of multiple stressors driven by global changes and the intensification of agriculture (Phillips *et al.* 2024; Rillig *et al.* 2023). While agricultural activities are essential for the survival and development of our society, intensive agricultural practices produce severe disturbance to soil biological communities (Yang *et al.* 2021), altering their composition, function, and ultimately impacting whole ecosystems and global health (Banerjee & van der Heijden 2023). For example, tillage (Angon *et al.* 2023), synthetic chemicals (Ruuskanen *et al.* 2023), deforestation (Souza *et al.* 2023), climate change (Banerjee & van der Heijden 2023), and biological invasions (Gao *et al.* 2022) have been linked to negative effects on soil health, loss of biodiversity, or variation in soil microbiome composition and function. Indeed, microbial communities inhabiting soil are key components in maintaining ecosystem functions and stability, with direct impact on plant health and tolerance to environmental stressors (Hartmann & Six 2022). For example, the effects of fire on the soil microbiome have been linked to reduction in plant productivity (Revillini *et al.* 2022), and, habitat fragmentation, through soil microbiome-mediated effects, has been found to influence plant performance and resource allocation (Kiesewetter & Afkhami 2021). However, we still know little about the influence of soil disturbance on the plant microbiome assembly and stability.

The soil microbiome has a major influence on the structure of plant microbiomes, in particular on those associated with belowground compartments (Malacrinò & Bennett 2024; Mohamed *et al.* 2024; Trivedi *et al.* 2020). While root microbiomes can differ significantly from bulk soil communities (Trivedi *et al.* 2020), environmental factors like drought and increasing temperatures (Tiziani *et al.* 2022) can have profound effects on the structure and functioning of the plant-associated microbial communities. Plants can indeed exert a certain degree of selection on their microbiome through the exudation of primary and secondary metabolites (Koprivova & Kopriva 2022; Sasse *et al.* 2017). These changes can, in turn, improve the ability of plants to forage nutrients and tolerate biotic and abiotic stressors (Pantigoso *et al.* 2022; Rolfe *et al.* 2019). Yet, our understanding of how environmental disturbance can alter plant-microbe interactions, and the assembly of the plant microbiome is still limited. In particular, previous studies heavily focused on bacterial communities, while we know much less about the influence of soil disturbance on the root fungal communities. While bacteria often dominate studies on soil microbiomes, fungi contribute significantly to nutrient cycling, organic matter decomposition, and plant health (Bahram & Netherway 2022; Banerjee & van der Heijden 2023; Fierer *et al.* 2021). Previous research has indeed shown that different farming systems, with varying levels of soil disturbance, have different effects on the assembly of the root microbial networks (Banerjee *et al.* 2019). While we have some evidence showing that soil disturbance can disrupt the assembly of plant-associated fungal communities, our understanding of how fungal microbiomes respond to varying levels of soil disturbance, particularly in roots, remains limited.

The assembly of plant-associated microbial communities is influenced by different ecological processes, where both stochasticity and determinism play a role in shaping the final structure of microbiomes at different compartments. However, how this assembly process is altered under different disturbance regimes remains an open question. Under stress, selection by the host plant may play a stronger role in selecting microbial taxa (Arnault *et al.* 2023; Barnes *et al.* 2024), favoring beneficial organisms that enhance nutrient acquisition and stress tolerance. For example, beneficial fungi may contribute to the robustness of microbial networks by stabilizing plant-fungal interactions and suppressing pathogens (Wang *et al.* 2024b). On the other hand, soil disturbance could also disrupt plant-microbiome interactions, leading to destabilized networks (Wang *et al.* 2024a) and promoting stochastic assembly processes, such as drift and dispersal. Understanding the balance between selection and stochasticity, as well as the role of different fungal taxa and functional groups in network stability, is crucial for predicting the consequences of soil disturbance on plant-microbe interactions. At the same time, understanding how microbiomes assemble can provide a solution to mitigate the negative effects of disturbances.

In this study, we asked whether soil disturbance can influence the assembly and the network of interactions within the root-associated fungal communities. We assembled microcosms, inoculated them with soils from sites with varying disturbance levels (agricultural field, field margins, uncultivated field) while controlling for soil abiotic features, and used amplicon metagenomics to infer the assembly processes and network of interactions behind the structure of root microbiomes. As observed in our previous studies (Malacrinò *et al.* 2021; Malacrinò & Bennett 2024), we predicted an inoculum-specific signature on the root fungal microbiome, thus reflecting the effect of environmental disturbance on the soil microbiome. We then asked if disturbance influences the ecological processes behind the assembly of the root fungal microbiome and hypothesized that higher levels of disturbance might favor selection processes, for example by recruiting beneficial organisms, with little contribution from stochastic processes. We also hypothesized that the robustness of the root fungal microbiome network would be influenced by the level of disturbance, predicting a lower robustness in plants exposed to the microbial communities derived from the agricultural soil, and a higher robustness in plants grown on microcosms inoculated with soil from the uncultivated field. Finally, we asked whether the fungal taxa supporting the network robustness are specific to each level of disturbance, expecting to observe inoculum-specific fungal taxa involved in both the assembly and robustness of the microbiome network. Considering that we expected a higher influence of deterministic assembly processes, as a consequence of potential microbial recruitment by the host plant, we also predicted that those taxa playing a major role in network robustness would mostly be identified as beneficial fungi.

## Results

We tested our hypotheses growing tomato plants (*Solanum lycopersicum* L. variety Moneymaker) in microcosms hosting three different soil microbial communities known to vary in composition. This approach was similar to our previous work (Malacrinò & Bennett 2024), as we sampled soils with different levels of disturbance: an agricultural soil (collected in a field cropped with tomato), the margins of that field (uncultivated area at the border between the uncultivated and the cultivated field), and an uncultivated field (undisturbed for the past ∼30 □years). The idea is that the agricultural field experienced the highest disturbance due to the conventional agricultural practices, the uncultivated field would bear minimal disturbances, while the field margins would be at an intermediate level of disturbance between the agricultural and the uncultivated field. Microcosms were assembled by mixing all three soils, of which only one was not autoclaved, thus maintaining alive the biological community (see Methods). This allowed us to control for abiotic factors and focus on the effects driven by the soil microbiomes in the three groups.

Each soil inoculum had a specific effect on the structure of the root fungal community (F = 8.93, R^2^ = 0.19, p < 0.001, Fig. 1A), also confirmed by pairwise post-hoc contrasts (FDR-corrected p < 0.01 for all combinations). Indeed, looking at the taxonomic composition (Fig. 1B), samples from the agricultural soil treatment were clearly dominated by ASVs identified in the genus *Rhizophagus* (63.3%), those from the field margin had higher abundances of fungi from the genera *Cladosporium* (21.8%) and *Epicoccum* (19.4%), while samples from the uncultivated soil treatment had higher abundances of ASV identified as *Funneliformis* (27.7%) and *Acaulospora* (22.7%). It is important to note that, among the most abundant taxa (Fig. 1B), the root microbiome of all samples includes both potential pathogens (e.g. *Epicoccum, Fusarium*) and beneficial organisms (e.g., mycorrhizal fungi from the genera *Acaulospora, Funneliformis, Rhizophagus*). Additional pathogens, including *Alternaria alternata*, were also identified among the less abundant taxa. Interestingly, the ASVs identified as beneficial organisms were those with the highest discriminatory power among the three soils using both the Random Forest (Fig. 2A) and the LefSe (Fig. 2B) approaches.

**Figure 1.**
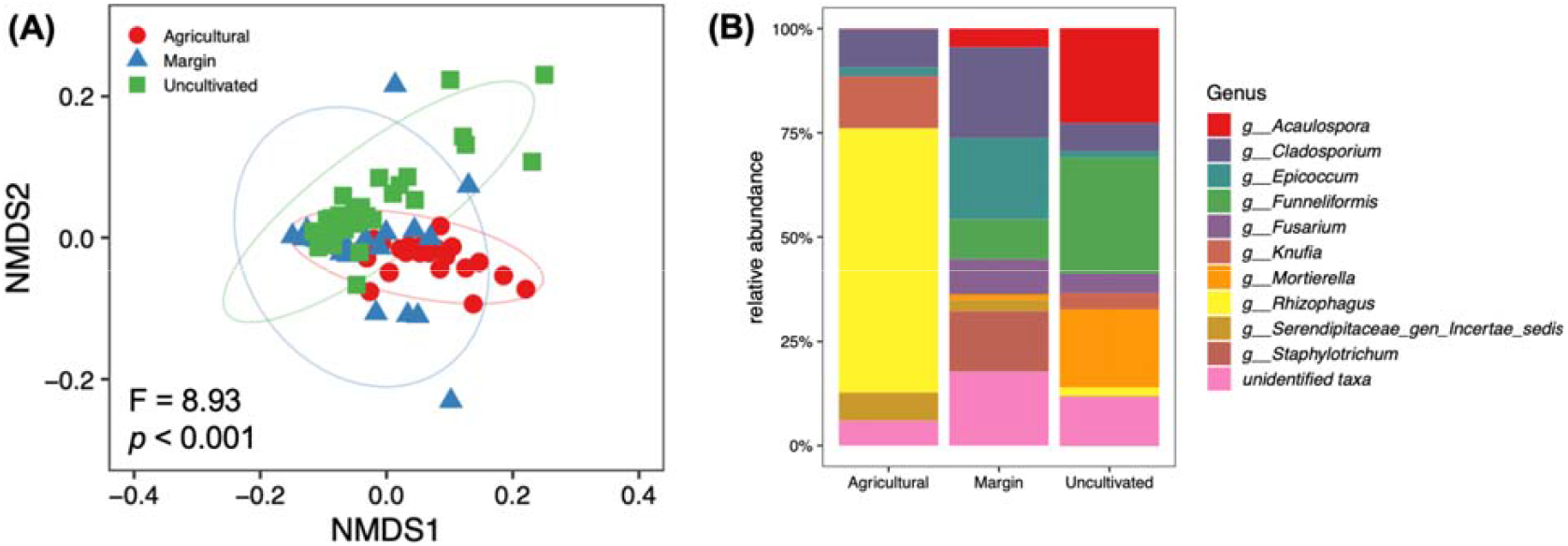
(A) Non-metric multidimensional scaling (NMDS) of the root fungal community built on a weighted UniFrac distance matrix, with points colored by soil inoculum. (B) Composition of reads of major fungal genera (relative abundance >1%) across soil inocula.

**Figure 2.**
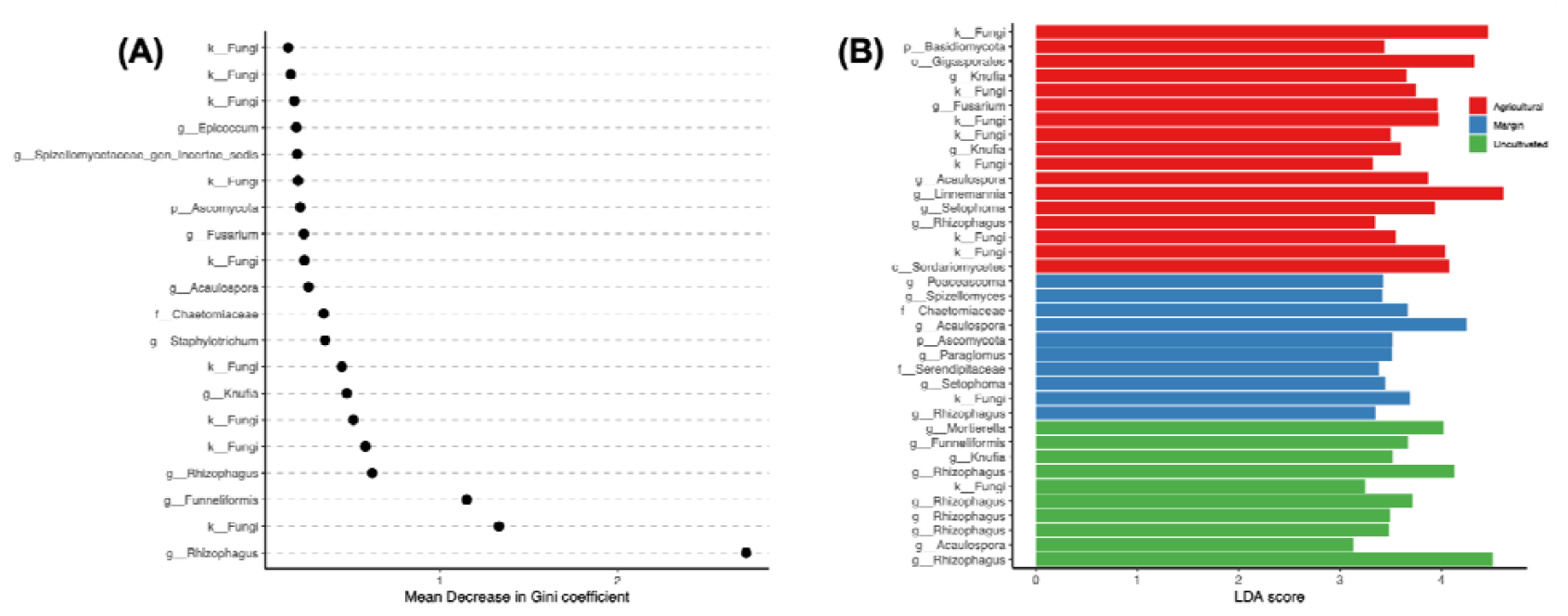
(A) Random forest analysis of the root mycobiome. The y-axis ranks the top 20 ASVs (identified at the highest confidence taxonomic level) ranked by their importance (Mean Decrease in Gini coefficient) for the group classification. (B) LEfSe bar chart showing the ASVs (identified at the highest confidence taxonomic level) with the strongest association to each of the three soil inocula, and the lengths of the bars (LDA score) indicate the influence of that ASV in discriminating between soils.

When comparing the diversity of the root fungal communities of plants grown on the different soils, we found no differences between the three groups using Shannon’s diversity index (Fig. 3A) and Faith’s phylogenetic diversity index (Fig. 3B) as metrics. However, when using the MNTD as a metric, we found a higher diversity in roots grown on the soil from field margin compared to the other two groups (Fig. 3C). We then compared the observed MNTD with a randomly assembled microbiome, using the β-NTI metric, and we found that all three groups had values between 2 and -2 (Fig. 3D), suggesting an assembly dominated by stochastic processes, which was then confirmed by further null-model analyses, showing that in all three groups drift and dispersal contributed more than selection in assembly the root fungal communities (Fig. 3E).

**Figure 3.**
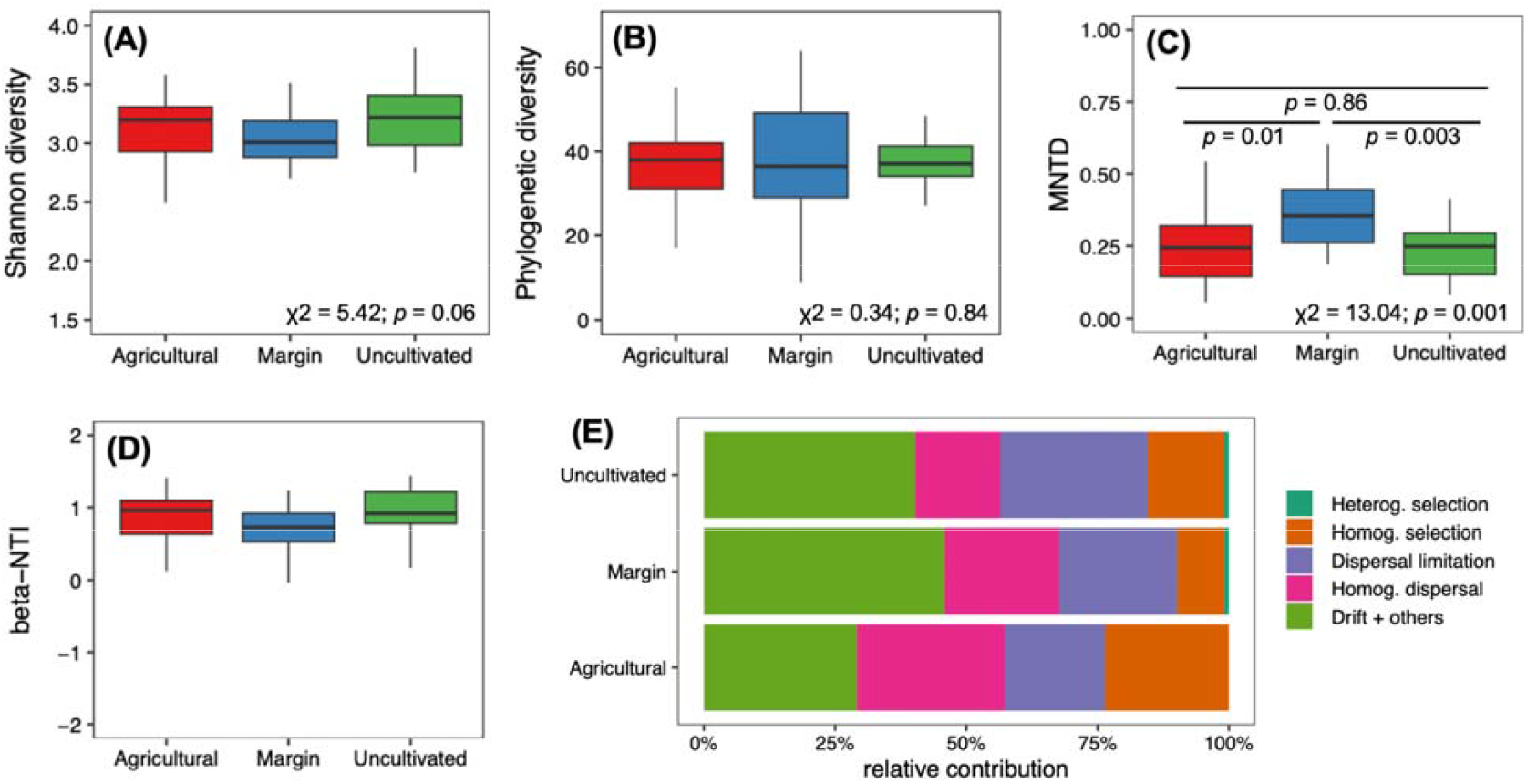
Comparison of Shannon’s diversity index (A), Faith’s phylogenetic diversity index (B), and Mean Nearest Taxon Diversity (MNTD) metric (C) of the fungal community in roots grown on the three different soils. For all three metrics, we show the results from a linear-mixed effects model and for the MNTD we show the FDR-corrected post-hoc contrasts. (D) β-NTI metric shows values between 2 and -2 suggesting an assembly similar to a null-model. (E) Null-model analysis showing the relative contribution of different processes in the assembly of root fungal communities in the three soils.

We then built a correlation network between taxa for each soil inoculum (Fig. 4A-C, Tab. 1), showing that the network complexity increased according to the level of disturbance, from agricultural soil, to field margin, and to uncultivated field. Indeed several metrics (Tab. 1) followed this pattern, including the number of nodes, the average path length, and the network modularity. Similarly, we observed an opposite trend for other metrics, showing that decreasing disturbance generated a less dense and centralized network (Tab. 1).

**Table 1.** Network metrics for each soil inoculum.

| Metric | Agricultural | Margin | Uncultivated |
| --- | --- | --- | --- |
| *Nodes* | 64 | 103 | 120 |
| *Edge* | 218 | 319 | 315 |
| *Average degree* | 6.81 | 6.19 | 5.25 |
| *Average path length* | 1.83 | 2.51 | 2.99 |
| *Network diameter* | 6 | 6 | 8 |
| *Clustering coefficient* | 0.71 | 0.60 | 0.61 |
| *Density* | 0.11 | 0.06 | 0.04 |
| *Heterogeneity* | 1.04 | 0.88 | 0.80 |
| *Centralization* | 0.25 | 0.12 | 0.09 |
| *Modularity* | 0.27 | 0.58 | 0.69 |

**Figure 4.**
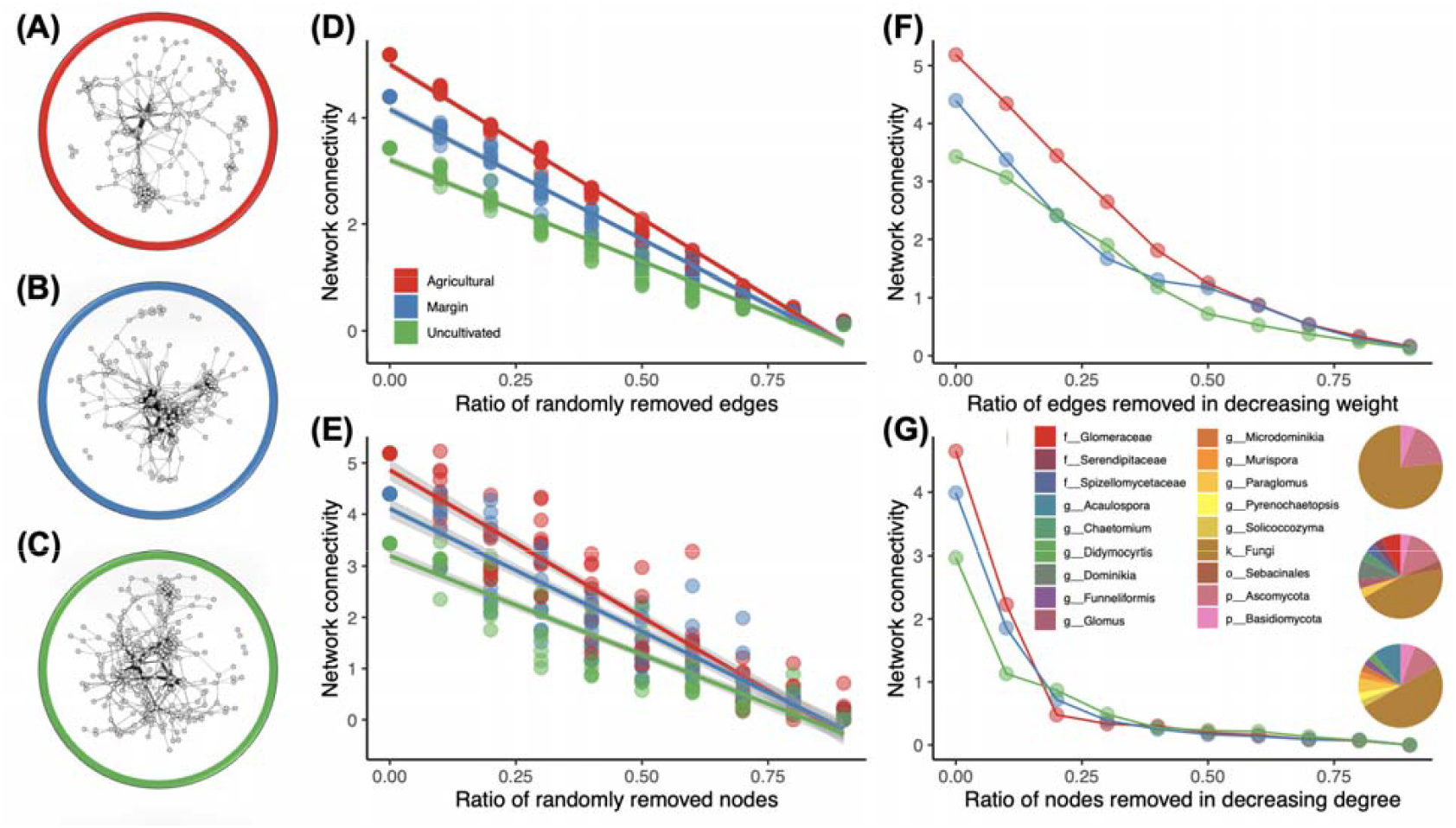
Network of fungal microbiome correlations for (A) agricultural field, (B) field margin, and (C) uncultivated field. Network connectivity as measure of network robustness as response to removing random edges (D) or nodes (E), and to removing edges in decreasing order of weight (F) or nodes in decreasing degree (G). Inset of panel G shows the taxonomic classification of the top 25% nodes for each soil group: agricultural field (top), field margin (middle), and uncultivated field (bottom).

Considering that several network metrics followed the disturbance gradient, we inferred the networks to test their robustness, and identify which taxa might best contribute to it. We first tested it by randomly removing edges (Fig. 4D) or nodes (Fig. 4E), and found that the network connectivity linearly decreased with their removal, with higher slopes for plants grown on agricultural soil, followed by field margin, and uncultivated field (Fig. 4D-E). We also tested the network robustness by removing edges and nodes for their relative importance within the network, rather than by random choice. When removing edges by decreasing weight (Fig. 4F) we observed a similar trend of random removal (Fig. 4D). However, removing nodes by decreasing degree showed that root fungal communities in plants grown on agricultural soil were much less robust than field margins or the uncultivated field, with the network connectivity sharply decreasing when removing <25% of most important nodes (Fig. 4G). We also found that the number of nodes with high degree (top 25%) was increasing from agricultural soil (n = 17), field margin (n = 27), and uncultivated field (n = 36) and, interestingly, in the agricultural soil none of those ASVs was identified to the species level while, for other two groups, several nodes were identified as mycorrhizal fungi (e.g., *Acaulospora, Funneliformis, Glomus, Paraglomus*, Fig. 4G).

The results above suggest that in our experiment the root fungal microbiome was assembled mainly following stochastic processes, however the network analysis suggested that some taxa might instead be key to network robustness and might not follow the same rules. We thus fit the data from each soil group to a Sloan null model (Sloan *et al.* 2006), which differentiated taxa following a null-model assembly from those that did not assemble according to a predicted random assembly (Fig. 5A-C). Then, we compared those taxa falling outside the predicted assembly and found that ∼14% were shared among the three soils (Fig. 5B), while the widest portion was unique to each soil inoculum. Interestingly, within the shared portion, no ASV was identified beyond the phylum level, while several ASVs unique to each group were identified as mycorrhizal fungi. In addition, for each soil inoculum we tested whether the ASVs falling outside a null-model assembly were the same as the top 25% high degree nodes as identified above, and we found that both the overlap (20%, 27.7%, and 29.8% for agricultural, margin, and uncultivated soil) and the diversity of fungal ASVs inside that overlap were increasing when reducing the disturbance level (Fig. 5C). Also in this case, mycorrhizal fungi (e.g. Glomeraceae, *Acaulospora*, and *Microdominikia*) were mainly represented in less disturbed soil (margin or uncultivated inoculum) compared to the one from the agricultural field (Fig. 5C).

**Figure 5.**
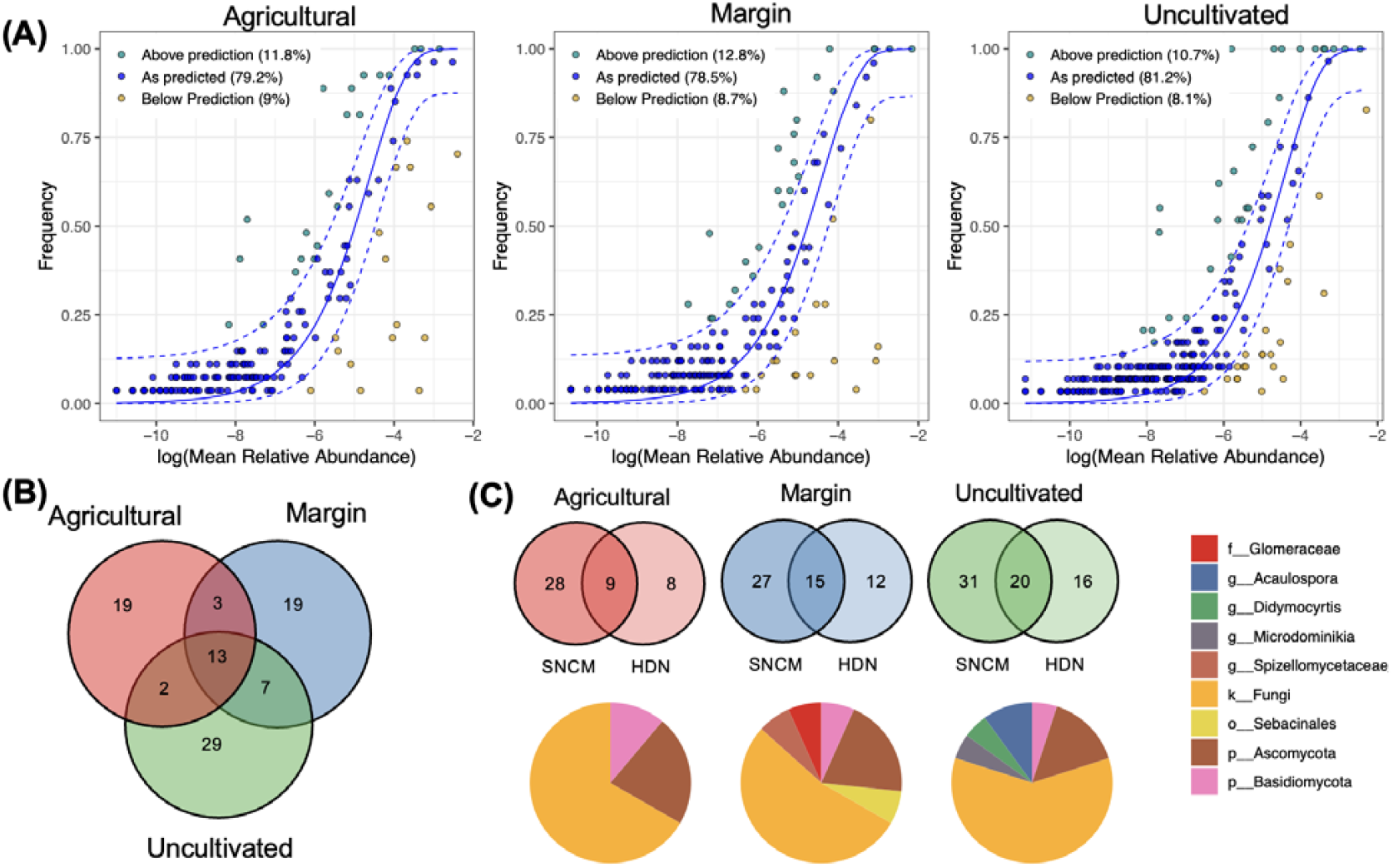
(A) Fit of root fungal microbial community ASVs to the Sloan null model for plants grown on microcosms inoculated with soil from the agricultural field (left), field margin (center), and uncultivated field (right). (B) Overlap of ASVs not fitting the Sloan null model between the three inocula. (C) Overlap of ASVs not fitting the Sloan null model (SNCM) with the high-degree nodes (HDN), for each soil type, and taxonomic classification (pie charts) for the overlapping ASVs.

## Discussion

In this study, we found that soil disturbance influences the assembly and robustness of root fungal microbiomes. Each soil inoculum produced a distinct root fungal community structure. In addition, we found that the robustness of the root fungal microbiome network (measured as variation in connectivity) decreased along the disturbance gradient, with the most complex and robust networks observed in plants grown on soil from the uncultivated field. Interestingly, we found that arbuscular mycorrhizal fungi (AMF) were major drivers in microbiome assembly and robustness. Furthermore, our results suggest that stochastic processes, such as dispersal and ecological drift, played a dominant role in shaping fungal community assembly across all soil treatments, although key taxa involved in network robustness did not follow a null-model assembly process.

Our results show that the source of soil inoculum had a strong influence on the structure of root fungal communities. Several studies reported that the origin of soil inoculum is a major driver in structuring plant-associated bacterial (Beschoren da Costa *et al.* 2022; Hannula *et al.* 2019; Malacrinò *et al.* 2021; Malacrinò & Bennett 2024; Tkacz *et al.* 2020; Zarraonaindia *et al.* 2015) and fungal communities (Almario *et al.* 2017; Beschoren da Costa *et al.* 2022; Bonito *et al.* 2019; Hannula *et al.* 2021). Soil origin, similarly to geographic location, might simply determine the initial pool of microorganisms available for root colonization, from which the plant can assemble their own microbiome. While we would have expected differences in fungal diversity between soils with different levels of disturbance, we did not observe major changes. Previous studies reported that not all the soil disturbance events influence fungal communities (Beschoren da Costa *et al.* 2022; Rodriguez-Ramos *et al.* 2021; Vahter *et al.* 2022), and no differences in fungal diversity were observed in root fungal communities across samples from farms with different levels of agricultural intensity (Banerjee *et al.* 2019). However, when looking at the MNTD index, we observed a higher diversity in soil from field margin compared to the other two groups. While it is tempting to link this observation to the intermediate disturbance hypothesis (Santillan *et al.* 2019), which posits an increase in biological diversity at intermediate levels of disturbance, this result might be simply due to the site-specific microbiome dynamics and further investigations are needed to confirm this result. However, if true, this result would confirm field margins as gatekeeper of biodiversity, as observed in for other biological communities (Crowther *et al.* 2023; Pérez-Méndez *et al.* 2025; Plath *et al.* 2021), supporting their importance in safeguarding also the microbial biodiversity in agricultural systems. Interestingly, we also observed that the root fungal microbiome in our system assembles, mostly, by following stochastic processes. This might be due to the fact that plants were exposed to the inocula for only 6 weeks and fungal communities respond differently from prokaryotic communities (Liu *et al.* 2024; Romdhane *et al.* 2022), so that their response to selective forces might need more time to be actually observed, as we noted in our previous short-term study (Mohamed *et al.* 2024). However, as discussed below, some taxa did not follow stochastic assembly processes.

Environmental disturbance has been previously linked to variation in soil microbiome network properties (Hernandez *et al.* 2021; Ortiz-Álvarez *et al.* 2021). In this study, we report for the first time the major role of AMF in the assembly processes and network robustness of the plant root microbiome, bridging and expanding the results from two previous studies. Zandt et al. (2023) used long-term mesocosm experiment established using soil with minimal disturbance (natural grassland and soil uncultivated for 60 years) to study the relationship between plant community stability and soil microbial networks. While they focused on soil rather than roots, the study observed a reduction in network stability in soil that experienced agriculture, but the authors did not link the effects they observed to any specific taxonomic or functional group. Banerjee et al. (2019), instead, focused on the root fungal microbiome of samples collected in farms performing agriculture at different intensities (conventional, no-till, and organic farming). Similarly to our study, they observed that less intense agriculture has the most robust fungal network, and that AMF are keystone species within these networks. In our experiment, we consider a wider range of disturbance compared to the individual previous studies, and we also show two interesting novel aspects. First, we assessed network robustness by measuring network connectivity after removing nodes/edges randomly or by their order of importance within the network. This allowed us to observe that networks from more disturbed systems are less robust, especially to the removal of high-degree nodes, which were mostly identified as AMF. Second, by null-model analysis, we observed that ∼25% of those high-degree nodes are also taxa that, differently from the rest of the microbiome, do not follow stochastic assembly processes. Taken together, these results provide strong evidence of the role of AMF in structuring the root-associated fungal communities and in supporting the robustness of such microbial networks. This is in agreement with the recognized role of AMF in plant ecology and evolution, ecosystem engineers, and gatekeepers of soil health (Carrara *et al.* 2023; Zhang *et al.* 2024). Indeed, plant-AMF symbiosis has important consequences for the host, including resistance to stressors, productivity, and overall interactions with other members of the surrounding biological community above- and below-ground (Dey & Ghosh 2022; Moora *et al.* 2025; Müller 2021; Sharifi & Ryu 2021; Shi *et al.* 2023; Vandenkoornhuyse *et al.* 2015).

Interestingly, in addition to AMF, several ASVs that played a key role in network robustness, and did not follow null assembly processes, were not identified beyond the phylum level. Also, this was more common in soil from the agricultural field and its margins compared to the uncultivated field. Often reference databases used for the taxonomic identification of ASVs, UNITE in our case (Abarenkov *et al.* 2024), do not include the full available knowledge on environmental fungal isolates because publicly available sequences – the main source for such databases – are often poorly annotated (Abarenkov *et al.* 2022), reducing our ability to correctly identify the full fungal diversity in a given sample using metagenomics tools. While this might partially explain our observations, we also have to acknowledge that much of the soil biodiversity is still uncovered and many of these microbial taxa might have never been cultured, be unculturable, or difficult to cultivate in vitro (Lewis *et al.* 2021). However, this less known portion of the soil and plant microbiomes, often called Microbial Dark Matter, might have key functional roles with important consequences for plants (Jiao *et al.* 2021). This provides support to the shift towards the use of more in-depth techniques, such as shotgun metagenomics and metatranscriptomics (Li *et al.* 2023), which is much needed and it might revolutionize how we understand plant-microbiome interactions in the wider framework of holobiont biology (Bordenstein *et al.* 2024).

Although we controlled for possible differences in soil chemistry and structure, a wider set of drivers, including the site-specific abiotic environment, might have exerted strong selection forces on the soil microbiome and, in turn, on the recruitment of fungal associates in our microcosms. In addition, while our study provides similar results using different approaches (network analysis, machine learning, and null-model analysis), further experimental evidence is needed to disentangle the role of AMF and other fungal taxa in the assembly of the root microbiome beyond the fungal community, additionally focusing on prokaryotes, nematodes, and other component of the soil- and plant-associated microbial communities. Research in this direction is essential to fully dissect plant-microbiome interactions and deploy microbiome-based strategies to restore and maintain soil and planetary one health (Banerjee & van der Heijden 2023; Compant *et al.* 2025; Montgomery *et al.* 2024; Singh *et al.* 2023; Yiallouris *et al.* 2024). This is especially important as the resilience of plant communities to the global changes we are facing, and are going to face, much depends on how such changes will impact plant-microbiome interactions (Addison *et al.* 2024; Trivedi *et al.* 2022), in particular considering that AMF might be more sensitive to global warming than expected (Xu *et al.* 2022).

Overall, our findings provide strong support for the idea that soil disturbance disrupts fungal microbiome assembly and weakens microbiome network robustness in plant roots. By demonstrating that beneficial fungi play a central role in stabilizing root microbiomes, our study highlights the importance of maintaining soil biodiversity to promote plant health and ecosystem resilience. This is particularly important considering that plant-microbe interactions in roots and rhizosphere play an essential role in the plant holobiont health and productivity (Bai *et al.* 2022; Beattie *et al.* 2024; Berg *et al.* 2024; Sánchez-Cañizares *et al.* 2017; Vandenkoornhuyse *et al.* 2015). Strategies that promote beneficial fungal communities, such as reduced tillage, organic amendments, and diversified cropping systems, may help mitigate the negative effects of soil disturbance and enhance microbiome resilience. Our findings further highlight the critical role of AMF in maintaining stable root microbiomes and suggest that promoting mycorrhizal diversity may be a key strategy for enhancing the resilience of plant communities in the face of increasing environmental disturbance. Management practices aimed at promoting AMF diversity, such as reduced tillage and cover cropping, could enhance the resilience of agricultural systems to disturbance and reduce reliance on external inputs. Future research should also investigate the functional roles of less known key fungal taxa and explore how these interactions influence plant health under different environmental conditions, further contributing to developing sustainable agricultural practices that harness microbial diversity to improve ecosystem stability and agricultural sustainability.

## Materials and methods

### Microcosm experiment

We tested our hypotheses by assembling microcosms using soil from sites with different levels of disturbance: an agricultural soil (collected in a field cropped with tomato), the margins of that field (uncultivated area at the border between the uncultivated and the cultivated field), and an uncultivated field (undisturbed for the past ∼30 □years). At each site, soil was collected between 5-15□cm of depth, sieved to 2□cm to remove large debris, homogenized, and stored at 4□°C. Sterilized background soil was generated from a mix of all three soils used for inoculation (1:1:1), which were then mixed with the same volume of sand (1 part combined soil: 1 parts sand). This mixture was sterilized by autoclaving at 121□°C for 3□h, allowing it to cool for 24□h, and then autoclaved at 121□°C for a further 3□h.

Microcosms were assembled in three groups, one of each inoculum sampled above. Pots (1L) were filled with a 1:1 (v/v) mix of sand and a soil mix. Sand was autoclaved at 121□°C for 3□h, allowing it to cool for 24□h, and then autoclaved at 121□°C for a further 3□h. The soil mix was generated from a mix of all three soils used for inoculation (1:1:1), of which only one was not autoclaved. For example, in the group of microcosms with the alive inoculum from the agricultural soil, each pot received a mix of 450mL of autoclaved sand and a soil mix of 150mL of alive (non-autoclaved) soil from the agricultural field, 150mL of autoclaved soil from the field margin, and 150mL of autoclaved soil from the uncultivated field. Using this approach, soil physicochemical properties were homogenized across treatments to isolate microbial effects from abiotic variation. Each group of microcosms was replicated over 32 pots, for a total of 96 pots divided into 4 blocks (24 pots of each inoculum per block). Pots were fully randomized between blocks and within each block, accounting for spatial variability of the abiotic environment within the greenhouse. A single tomato seedling (*Solanum lycopersicum* L. variety Moneymaker, previously germinated on autoclaved potting soil) was transplanted to each pot. Plants were then grown in a greenhouse at an average temperature of 25□°C and a photoperiod of 16□h of light and 8 h of darkness. Plants were watered with ∼100□mL of tap water three times per week throughout the experiment. Six weeks after the experimental setup, from each pot, we collected ∼50□mg of roots from each plant, carefully washed them with tap water, and immediately stored samples at -80°C.

### DNA extraction, library preparation, sequencing

Each sample was lysed in an extraction buffer using a bead-mill homogenizer, and total DNA was extracted using a phenol-chloroform protocol. We did not surface sterilize samples, so we characterized both endophyte and epiphyte communities. After quality check, we prepared libraries targeting the fungal ITS region of the rRNA using the primer pair ITS1-ITS2. Amplifications were also carried out on non-template controls, where the sample was replaced with nuclease-free water to account for possible contamination of instruments, reagents, and consumables used for DNA extraction. After this first PCR, samples were purified (Agencourt AMPure XP kit, Beckman Coulter) and used for a second short-run PCR to ligate Illumina adaptors. Libraries were then purified again, quantified using a Qubit spectrophotometer (Thermo Fisher Scientific Inc.), normalized using nuclease-free water, pooled together, and sequenced on an Illumina NovaSeq 6000 SP 250PE flow cell at the Genomic Sciences Laboratory of North Carolina State University (Raleigh, NC, USA).

### Data processing and analysis

Reads were processed with the *nf-core/ampliseq* v2.7.1 pipeline (Di Tommaso *et al.* 2017; Ewels *et al.* 2020; Straub *et al.* 2020) for quality control, adaptor/primer trimming, error correction, clustering of ASVs, chimera removal, and taxonomic identification using the UNITE database v10.0 (Abarenkov *et al.* 2024). ASV sequences were aligned using *MAFFT* v7.525 (Katoh *et al.* 2002) and used to build a phylogenetic tree using *FastTree* v2.1.11 (Price *et al.* 2009). All subsequent data processing and analysis was performed in *R* v4.4.1 (R Core Team 2020). The ASV table, taxonomic identification table, metadata, and phylogenetic tree were merged using *phyloseq* v1.48 (McMurdie & Holmes 2013) before subsequent analyses. Likely contaminants were removed using the package *decontam* v1.24 (Davis *et al.* 2018) and the data from negative control PCRs. Then, singletons and samples with less than 1,000 reads were discarded from the dataset. This resulted in a small reduction of replicates for each group (n = 27 for agricultural soil, n = 25 for field margin, n = 29 for uncultivated soil), but still maintaining a high level of replication, and a depth of ∼4,255 reads/sample, which clustered into 409 ASVs. Data used for calculating relative abundances were normalized using *Wrench* v1.22 (Kumar *et al.* 2018) before processing.

Differences in microbiome structure were tested using PERMANOVA (999 permutations stratified at the “block” level) and visualized using a NMDS, both performed on a weighted UniFrac distance matrix using the package *vegan* v2.6 (Dixon 2003). PERMANOVA post-hoc contrasts were performed using the package *RVAideMemoire* v0.9 (Hervé 2022). The random forest classification was performed using the package *randomForest* v4.7 (Breiman *et al.* 2002), while the LEfSE (Linear discriminant analysis Effect Size) analysis (Segata *et al.* 2011) was performed using the package *microeco* v1.13 (Liu *et al.* 2021). The Shannon diversity index was calculated using the package *microbiome* v1.26 (https://github.com/microbiome/microbiome/), the Faith’s phylogenetic diversity, and the MNTD and β-NTI metrics were calculated using the package *picante* v1.8.2 (Kembel *et al.* 2010). For each univariate metrics, differences between groups were tested by fitting a linear mixed-effect model using the package *lme4* v1.1 (Bates *et al.* 2015) using the variable “soil inoculum” as fixed factor and the variable “block” as random effect, while post-hoc contrasts were extracted using the package *emmeans* v1.10.7 (Lenth 2022). The relative contribution of different ecological processes in the assembly of the root fungal microbiomes for each soil inoculum was estimated using the package *iCAMP* v1.5.12 (Ning *et al.* 2020). Networks were built and analyzed using the package *microeco* v1.13 (Liu *et al.* 2021), and visualized using the package *igraph* v2.1.4 (Csárdi *et al.* 2025). Fit to the Sloan null-model was performed for each soil inoculum using the package *tyRa* v0.1 (https://github.com/DanielSprockett/tyRa).

## Acknowledgements

We acknowledge the support by Clemson University HPC resources (Antao *et al.* 2024).

## Author contributions

Conceptualization: SM, AM; Methodology: SM, LS, AM; Investigation: SM, EF, MMA, NZM; Visualization: SM, AM; Funding acquisition: AM, LS; Writing-original draft: SM, AM; Writing-review and editing: all co-authors.

## Data and code availability

Data is available on NCBI SRA under BioProject PRJNA1256102. All code to replicate data analysis is available at: https://github.com/amalacrino/mosca_et_al_RootFungalNetwork/.

